# MOAST: Mechanism of Action Similarity Tool

**DOI:** 10.1101/2025.09.15.676411

**Authors:** Akshar Lohith, Derfel Terciano, Adam Murray, John B. MacMillan, R. Scott Lokey

## Abstract

Determining the mechanism of action (MOA) for natural products remains a significant bottleneck in drug discovery, particularly for researchers with limited computational resources or small compound libraries. Traditional approaches require screening large numbers of annotated compounds alongside unknowns, which is cost-prohibitive, or depend on complex machine learning models that need substantial computational resources and large datasets.

Here, we present a dissertation chapter excerpt: MOAST (Mechanism of Action Similarity Tool), a BLAST-inspired computational workflow that addresses these limitations by providing rapid MOA hypotheses for newly screened compounds. This chapter investigates two complementary approaches: a kernel density estimation (KDE) method providing statistical significance measures and E-values for MOA class membership, and a CatBoost machine learning classifier for multi-class prediction with ranked outputs.

Using cytological profiling data from HeLa and A549 cell lines, MOAST achieved 22% accuracy for the top 5 predictions among ∼ **300** MOA classes, with the CatBoost classifier reaching 10% balanced accuracy—significantly better than the ∼ **3**% reported in literature. The tool suggests a 0.8 prediction probability threshold for trustworthy results and demonstrates robust performance across multiple feature reduction strategies.

MOAST provides a practical, accessible solution that bridges traditional phenotypic screening and modern computational approaches, making MOA determination feasible for researchers with limited resources while maintaining statistical rigor and interpretability.

## 1 Introduction

Natural products have been pervasive in human history as sources of drugs and medicines. Be it traditional herbal remedies or accidental discoveries, compounds extracted from or derived from natural sources have been crucial to the development and inspiration for over 1000 therapeutics in the last 30+ years. The traditional method used to uncover and isolate new compounds from natural sources is glibly called the “grind and find” method. These methods would start with observing therapeutic activity, followed by iterative, bioassay-driven, pure culture isolation, extract fractionation, and mass spectrometry to isolate and characterize a lead compound. Then, biological and biochemical studies characterize the compound’s functional mechanism and target (mode/mechanism of action).

With the innovation and widespread availability of automated instrumentation and techniques, more chemical products can be analyzed with high-throughput screening, screening of many lead compounds quickly in 96-well or greater format, and highcontent screening, collecting and analyzing multiple measurements from the same screen Schulze et al (2013). Phenotypic screening has emerged as a powerful approach for drug discovery, particularly for natural products and compounds with unknown mechanisms of action Moffat et al (2017). High-content imaging assays, such as cytological profiling (CP), generate rich phenotypic fingerprints that capture cellular responses to compound treatment Caicedo et al (2017). However, interpreting these complex multivariate signatures to determine the mechanism of action (MOA) remains a significant challenge.

Traditional approaches to MOA determination often rely on trained machine learning models that require substantial computational resources and large, well-annotated datasets. While these methods have produced exciting results and advanced drug discovery, they present barriers for researchers with limited resources or smaller compound libraries. The models themselves are not readily available to the public, and reproducing results requires building and training fresh models on similar data, where maintaining repositories of labeled data becomes challenging. Additionally, traditional screening studies require screening large numbers of annotated compounds alongside unknowns, which can be cost-prohibitive for many research groups.

An alternative approach involves analyzing the enrichment of MOA classes among the most highly correlated phenotypic fingerprints. This strategy inspired the development of MOAST (Mechanism of Action Similarity Tool). This BLAST-inspired analysis workflow provides MOA predictions for small sets of phenotypic fingerprints through a database query model Altschul et al (1990). MOAST takes a query compound fingerprint, computes its pairwise similarities to a set of known reference fingerprints, and produces likelihood probabilities for membership in known compound classes.

## 2 Methods

### 2.1 KDE Integration Method

For this BLAST-inspired analysis workflow, a sufficiently large database of phenotypic fingerprints is needed to produce a robust query mechanism that serves as the similarity analysis’s null model. A set of ‘random’ or artificial fingerprints would first need to be generated to build up the database’s size to have a null model. We first started by creating synthetic fingerprints, based on the reality of the ranges of HistDiff scores computed for each feature.

For every feature we have measured in the CP assay or any phenotypic fingerprint assay, we could draw distributions to help create artificial fingerprints that can serve as the null model (Supp. Figure 22). The artificial fingerprints are constructed by drawing scores from each of these distributions defined for each feature and composing them into the numerical vector that is the compound phenotypic HistDiff fingerprint. We tested multiple null model approaches, including unit normal distributions, feature-defined normal distributions, Gaussian kernel density estimation, and random sampling of annotated fingerprints with replacement.

With a database of artificial or randomly sampled fingerprints and computed similarity/distance scores to reference fingerprints, we can ask and answer how related a query fingerprint is to any MOA class represented in the reference set. The average correlation distance of reference fingerprints to the null model (artificial fingerprints or randomly sampled) can be grouped by reference compounds that share a specific MOA label and then used to compute a kernel density estimate (KDE) to serve as the label’s null model distribution.

Integrating a kernel density estimation (KDE) made from the average correlation distances of fingerprints annotated MOA class to the ‘random’ null model, from 0 to the average correlation distance of a fingerprint of an unknown/unlabeled fingerprint to that specific annotated MOA, and results in the cumulative distribution function (CDF) value corresponding to that average correlation distance. Specifically, the integral would be:

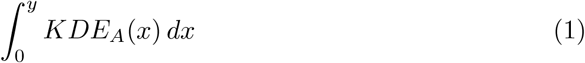

where *y* is the average correlation distance of the fingerprint of the unknown to fingerprints annotated for class A, and *KDE*_*A*_(*x*) is the kernel density estimation function constructed from the average correlation distances of fingerprints annotated for class A to a random collection of phenotypic fingerprints.

The calculation of the ‘avg corr dist unknown to A’ and evaluation of the KDE integral can then be repeated for each MOA class annotation, giving an array of independently calculated likelihood probabilities that the observed average correlation distance could have occurred by chance for the unknown to all the annotated MOA classes. Multiplying this KDE integrated probability by the number of MOA classes tested will result in an E-value, similar to what a BLAST search provides when returning results of the query sequence, allowing for a quick sort and interpretation of a potential relationship to an MOA class.

### 2.2 Conventional Machine Learning Classifier

Using the Python package ‘pycaret,’ multiple conventional machine learning methods can be tested together on the same data, with performance metrics measured and provided at the end in a tabular format. The primary metrics considered are accuracy, balanced accuracy, precision, and recall. The F1 score combines precision and recall in one value and is calculated as follows:

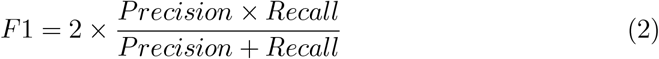

The F1 score ranges from 0 to 1, where 1 indicates perfect precision and recall, and 0 indicates the worst possible performance. As the chemical reference libraries are often imbalanced in terms of annotation representation, the F1 score is particularly useful to balance the trade-off between precision and recall, providing a more informative measure than just the straight accuracy of trained models when dealing with uneven class distributions or when false positives and false negatives have different costs.

The best model from this multiple testing is then trained and cross-validated—a process of partitioning the data into batches of held samples and training samples, iteratively refining the model training to ensure that all the learned feature weights can accurately represent the possible partitions in the training data in a multi-class cognizant manner. With the trained model, putative MOA annotation classes of unknown samples can be predicted, and the probabilities of annotation matches returned to the user.

### 2.3 Feature Selection Strategies and Evaluation

Feature selection is critical for the success of both KDE integration and machine learning approaches. We implemented a data-driven approach to feature selection that resolves shortcomings in determining class separations while maintaining robust clustering for strong MOA classes. First, uninformative features with zero standard deviation across all perturbations were removed, and then we further reduced redundancy using the findCorrelation function in the R-package caret. This function examines all feature pairs with Pearson correlation coefficients greater than a set threshold (0.95 in our case for HeLa CP assay data and 0.8 for A549 CP assay data), then flags the member of each pair with the highest mean correlation to all other features for removal, reducing collinearity among selected features.

We also explored alternative methods for feature reduction, including Principal Component Analysis (PCA) and Uniform Manifold Approximation and Projection (UMAP). While these methods can produce more manageable feature dimensions, the feature descriptions lose their experimental legibility, having been abstracted into composite representations. This feature reduction workflow was implemented on cytological fingerprints for HeLa CP assay using the SelleckChem reference library, a commercial set of compounds with known MOA and with some bioactivity and clinical interest. We then used a two-sample, one-sided Kolmogorov–Smirnov (KS) test to determine whether in-class associations differed from out-of-class ones as a proxy for the information content available in the fingerprints with different feature reduction strategies. We used this approach in a paper, High-Throughput Functional Annotation of Natural Products by Integrated Activity Profiling Hight et al (2022)

## 3 Results and Discussion

### 3.1 Feature Selection Performance

The first consideration of this method is that information provided in the fingerprint needs to be ‘fixed’ such that the building of the MOA null models and subsequent runs of the MOAST method rely on the phenotypic assay to collect and report on the same measurements. To implement and test this method, we started with the 245 CP assay features that we had determined to be the fixed feature set for the HeLa CP assay. Breaking down to the successive steps in the data processing of the HeLa CP assay, the image analysis step, where image masks are generated, and the source fluorescent images are measured for a variety of morphological features (area, pixel intensity, nucleus size, etc.), can produce hundreds or thousands of measurements, many related to each other.

Originally, to provide meaningful interpretations of phenotypic fingerprints, technical reproducibility was used to prune the phenotypic CP assay fingerprint to a set of 248 features from 430 Woehrmann et al (2013). This was done by assaying a selection of compounds with replicates and then performing a linear regression analysis of the measurement features from each replicate after scoring with the HistDiff algorithm - features with an *R*^2^ *>* 0.6 were kept as part of a fixed feature set. This set of empirically determined features still had shortcomings when resolving the fingerprints of closely related compound classes, in that features can still be highly correlative, providing redundant information in the fingerprint.

We then applied a data-driven approach to feature selection to resolve some of those shortcomings in determining the class separations while maintaining robust clustering for strong MOA classes. First, uninformative features with zero standard deviation across all perturbations were removed, and then we further reduced redundancy using the findCorrelation function in the R-package caret. This function examines all feature pairs with Pearson correlation coefficients exceeding a specified threshold (0.95 in this case) and flags the member of each pair with the highest mean correlation to all other features for removal, thereby reducing collinearity among the selected features. The HD scores in these selected features will result in a final CP fingerprint for each treatment condition.

This feature reduction workflow was implemented on cytological fingerprints for HeLa CP assay using the SelleckChem reference library, a commercial set of compounds with known MOA and with some bioactivity and clinical interest. We then used a two-sample, one-sided Kolmogorov–Smirnov (KS) test to determine whether in-class associations differed from out-of-class ones. The original fixed set of features had fewer classes defined as significant compared to the data-driven approach. We used this approach in a paper, High-Throughput Functional Annotation of Natural Products by Integrated Activity Profiling Hight et al (2022).

However, a fixed set of features can provide cross-screen comparability without repeating the feature selection process. Determining a fixed set of features using the data-driven approach to present to the public can help non-specialists of the CP assay compare compounds from different screening sets and hypothesize the MOA of their compounds of interest.

The exact features measured in the CP assay with the image analysis step remain at 430 (for the HeLa CP assay). When comparing individual screens, the data-driven feature reduction ultimately selects a different assortment of features to be a part of the final represented fingerprint. Since the measured set of features remains constant, a new fixed set of features can be determined to recover the original feature set’s cross-screen comparability. Previous CP experiments can be partitioned into groups to generate a new fixed feature set, and the feature reduction process can be applied to each group. The common features selected from the feature reduction on each partition can make up the new fixed set of cytological features.

To confirm that this new fixed set of cytological features does not distort from the improvements in the feature reduction process, the Venn diagram 1 demonstrates that MOA class significance performs just as well as the feature reduction process. The ‘5HTR.antagonist’ and ‘Carbonic.Anhydrase’ mechanistic classes are considered significantly distinct with just the feature reduction process and instead the Aromatase IkB.IKK mechanistic classes are considered significantly distinct from the new frozen set of features. However, these classes all exist near the significance threshold of p¡0.01.

We also investigated alternative methods for feature reduction, such as Principal Component Analysis (PCA) and Uniform Manifold Approximation and Projection (UMAP). While the defined feature dimensions that capture the most information can be much more manageable, the feature descriptions lose their experimental legibility, having been abstracted away into composite representations. Interestingly, when validating the feature reduction methods for the A549 iteration of the CP assay, we can see in Figures 2 and 3 that using the data-driven collinearity reduction can produce similarly defined class distinctions, as determined by the KS test. Even though the number of features is much larger than the number of features generated by the composite reduction methods, being comparable to the composite method could provide an avenue for when even a marginal reduction in the number of features can be beneficial for downstream processes, like the KDE integration methodology of MOAST.

**Fig. 1.**
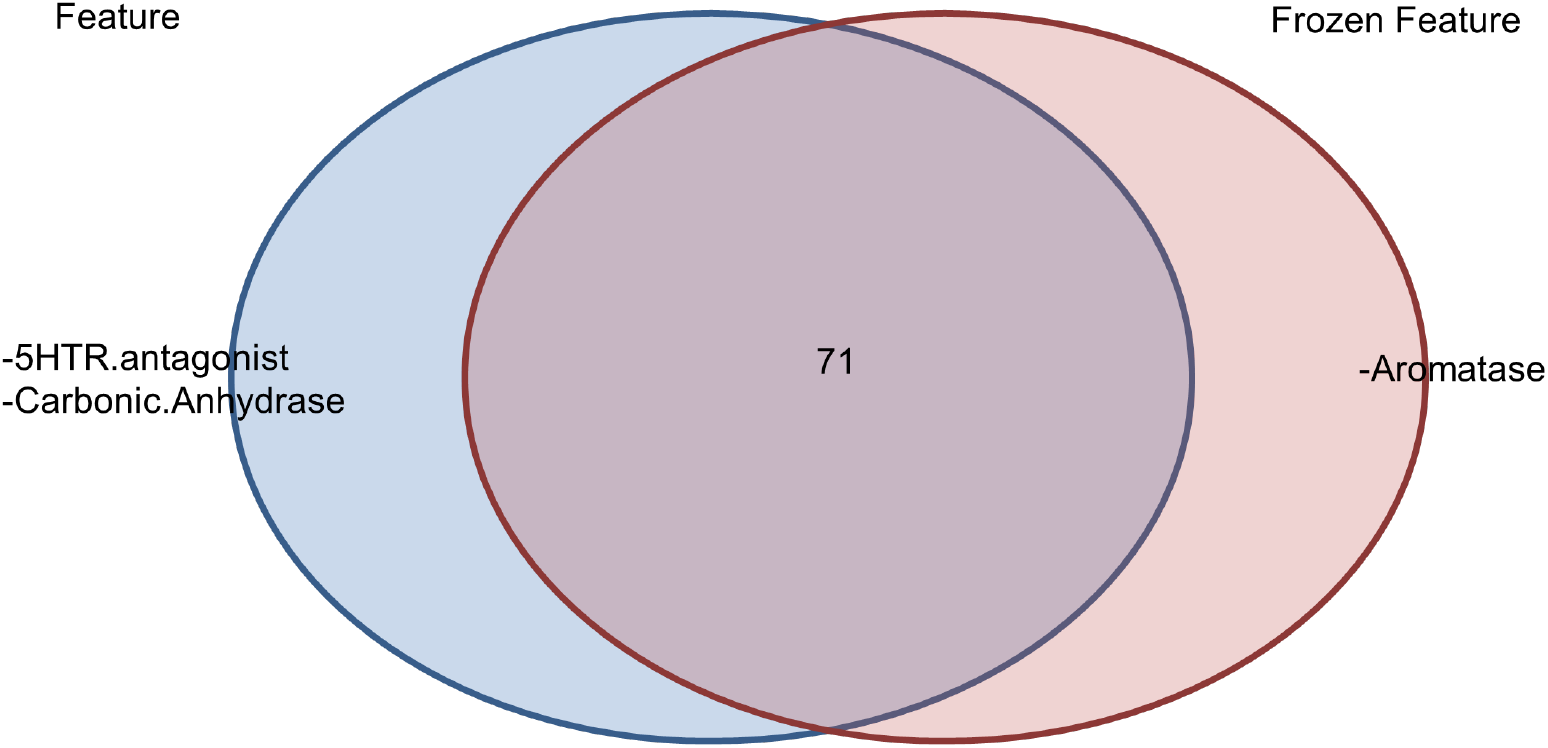
Venn-diagram demonstrates that MOA class significance performance with different feature sets, Empirical Features vs. the Feature Reduction process determined Frozen Feature set.

**Fig. 2.**
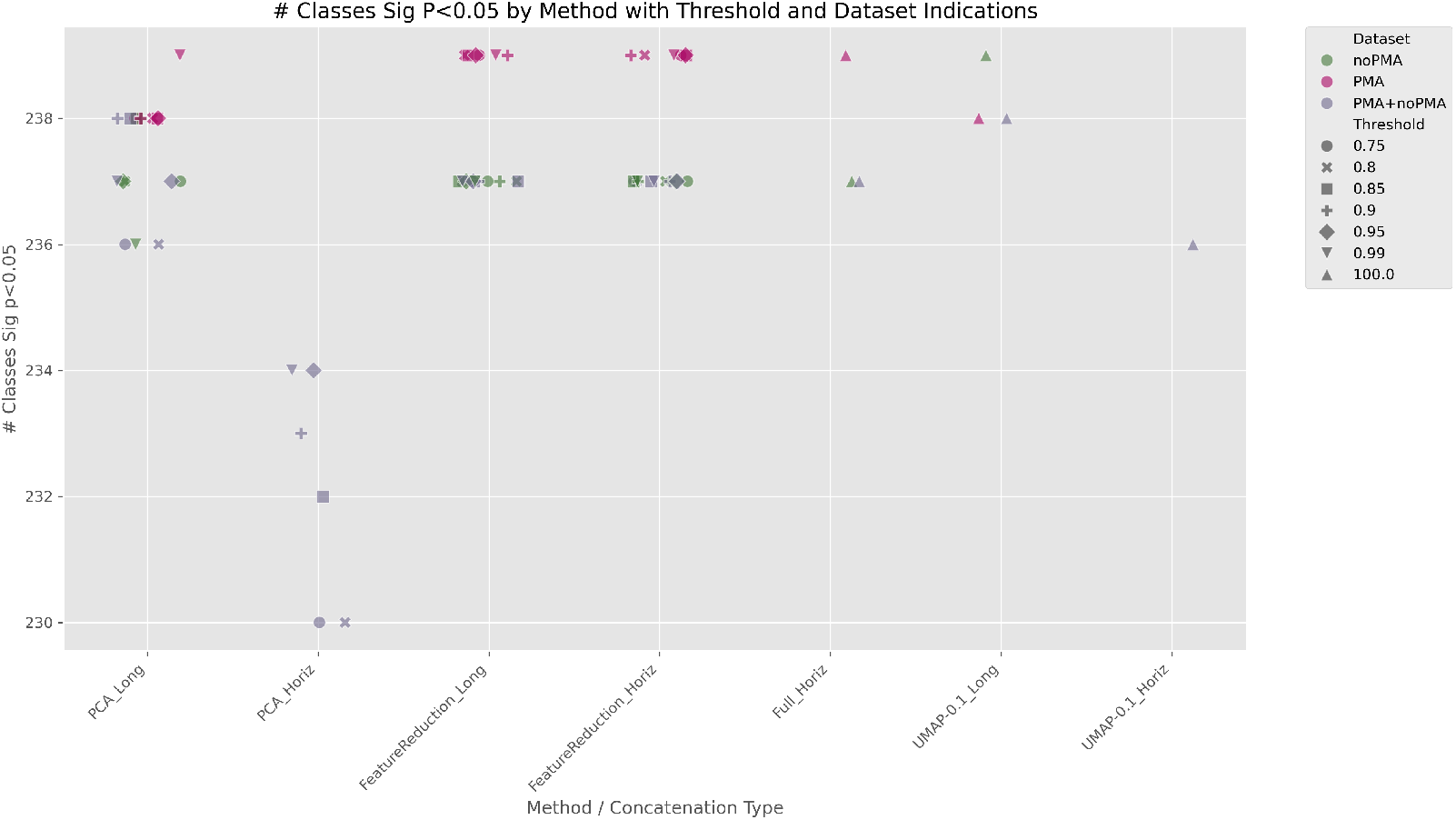
Number of classes significant (*p <* 0.05) following KS test of 241 annotated classes from A549 iteration of CP assay.

**Fig. 3.**
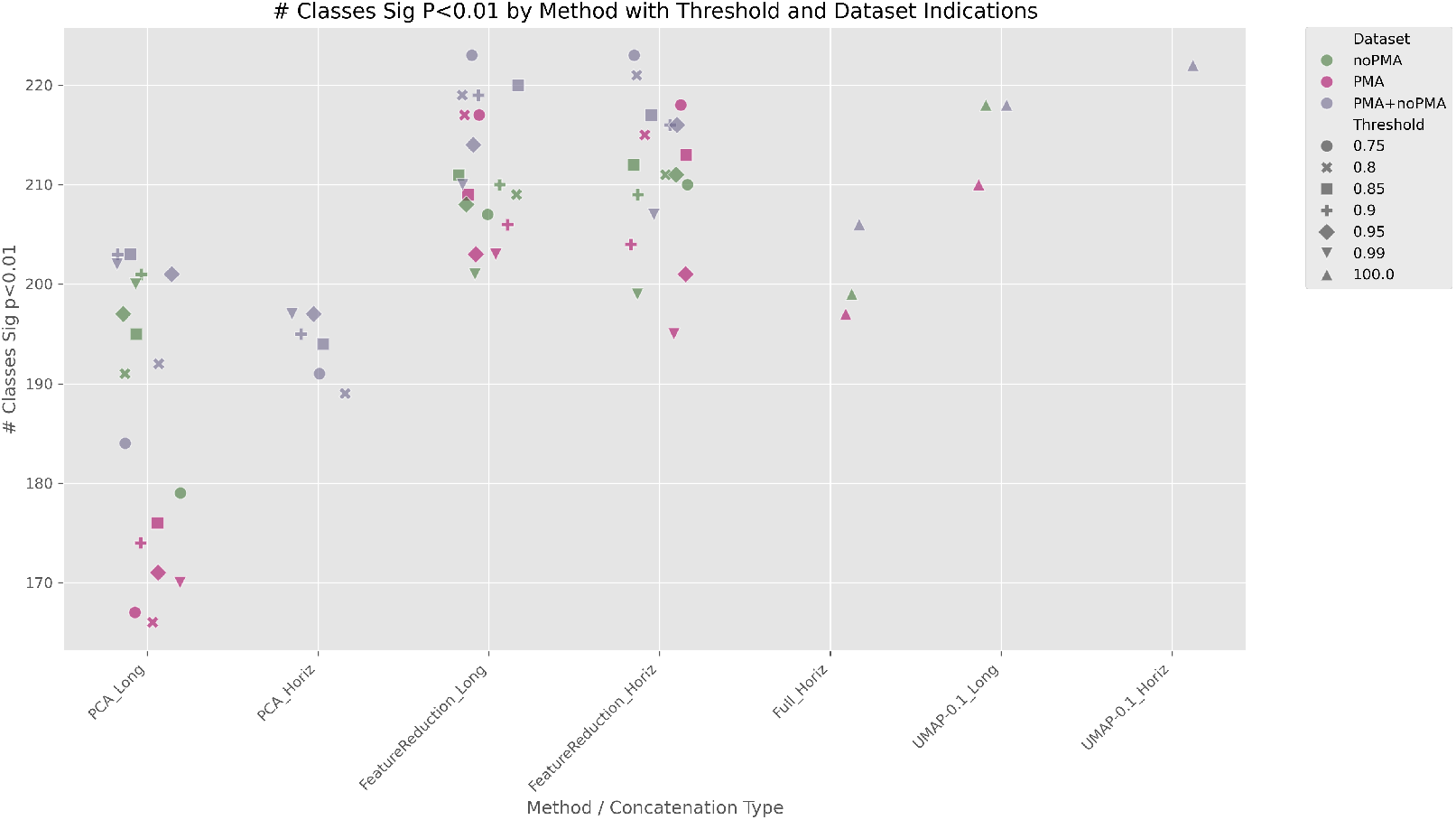
Number of classes significant (*p <* 0.01) following KS test of 241 annotated classes from A549 iteration of CP assay.

### 3.2 KDE Integration Validation

To validate this approach, we performed a 10% hold-out testing on the HeLa reference data, repeated ten times. This methodology goes like the following: a 10% sample of annotated compounds from the reference signatures is selected to be removed from the reference and held aside to serve as a set of unknown samples to be tested. Next, as defined in the method, the KDEs for each class represented in the modified reference set are constructed. Following this, each signature in the holdout is tested against all the classes, and the results are ranked based on the probabilities and proportions of when the held compounds’ annotation is among the top 1-5 ranked probabilities, which is calculated and represented for each tested null model below.

In figure 4, we demonstrate the ranking results when we only consider the ∼82 classes with enough representative membership to perform a KS test from the Hight et al. 2022 paper and pass a KS test significance threshold of p*<*0.01. As seen in this figure (4), the accuracy for the top class is relatively poor. It is probably primarily contributed to the interrelatedness of MOA annotations and the remaining noisy signal in the phenotypic signatures. The KDE integration methodology has a final reporting step of converting the P-values into E-values and then using the E-value to quickly rank and assess putative MOA annotations that could apply to the query phenotypic fingerprint for the assay. When looking at the distributions of the E-Values across ranks (5), a single E-Value that would apply to all classes and all ranks is difficult, or at least variable, depending on which rank is being considered. A systematic method of cutoff E-Values could be better than the median mismatch E-value for each rank, where matches are the E-Values for the held compound that have an MOA annotation that matches the corresponding MOA annotation. Mismatches are when the held compound has an MOA that does not match the corresponding MOA annotation. Alternatively, for each class, two KDEs can be constructed from the matches and mismatches, and the E-value of the intersection between the two can be the threshold E-value for the class.

**Fig. 4.**
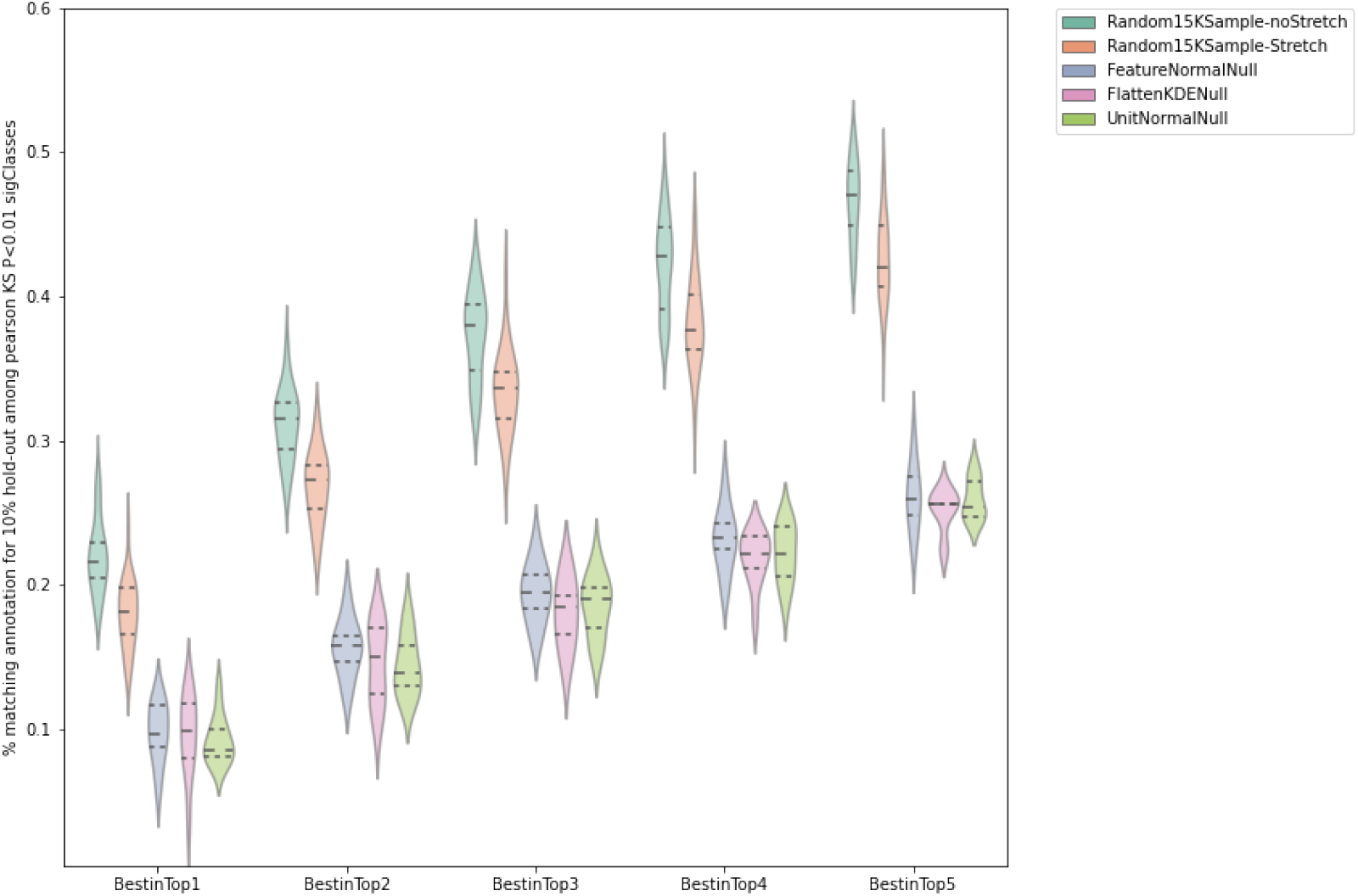
Accuracy of Top 1-5 across 10% held-out compound signatures for different tested Null Models.

**Fig. 5.**
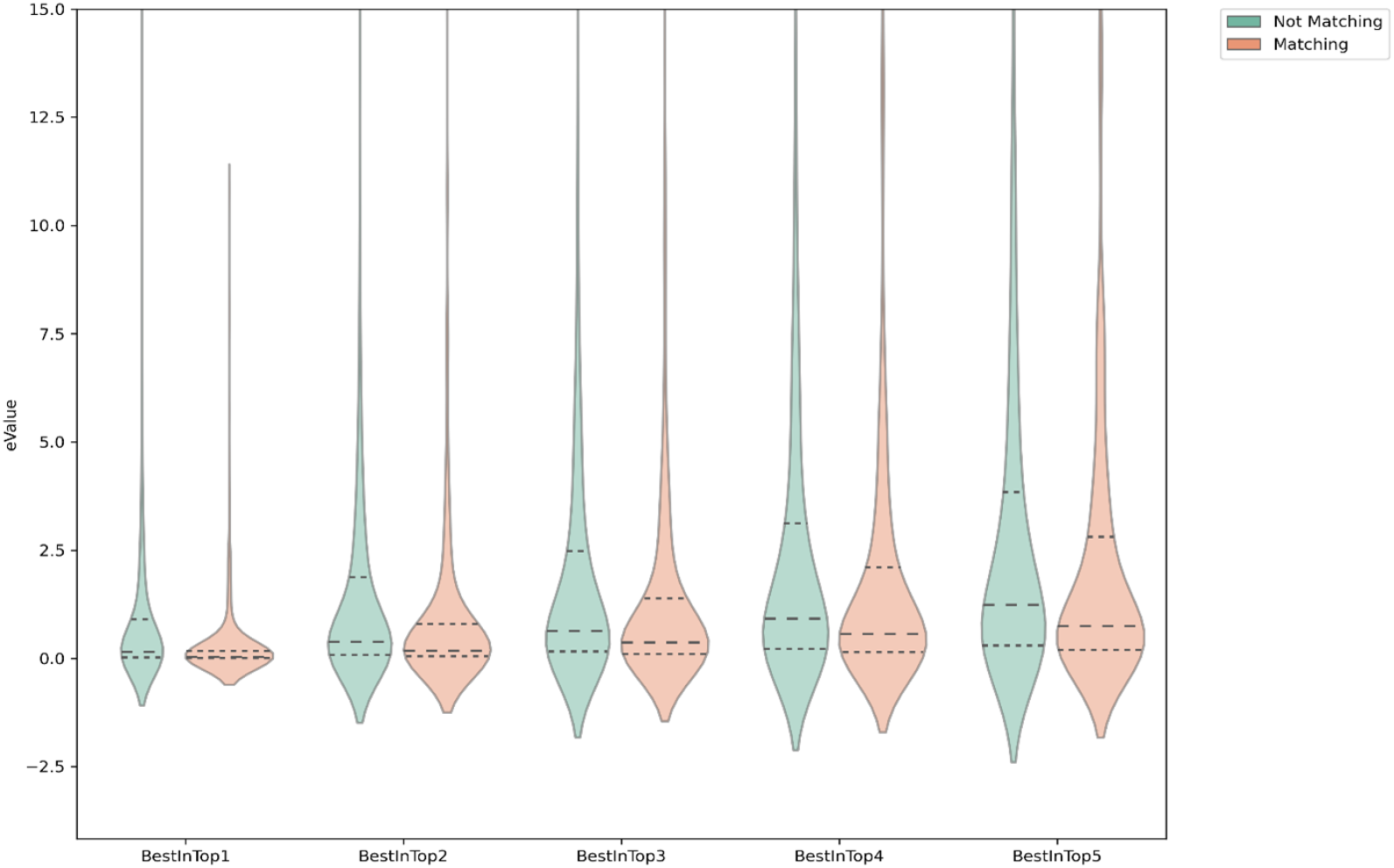
Distribution of E-Values for the Top 1-5 matching/mismatching annotations assigned by the KDE integration method.

**Fig. 6.**
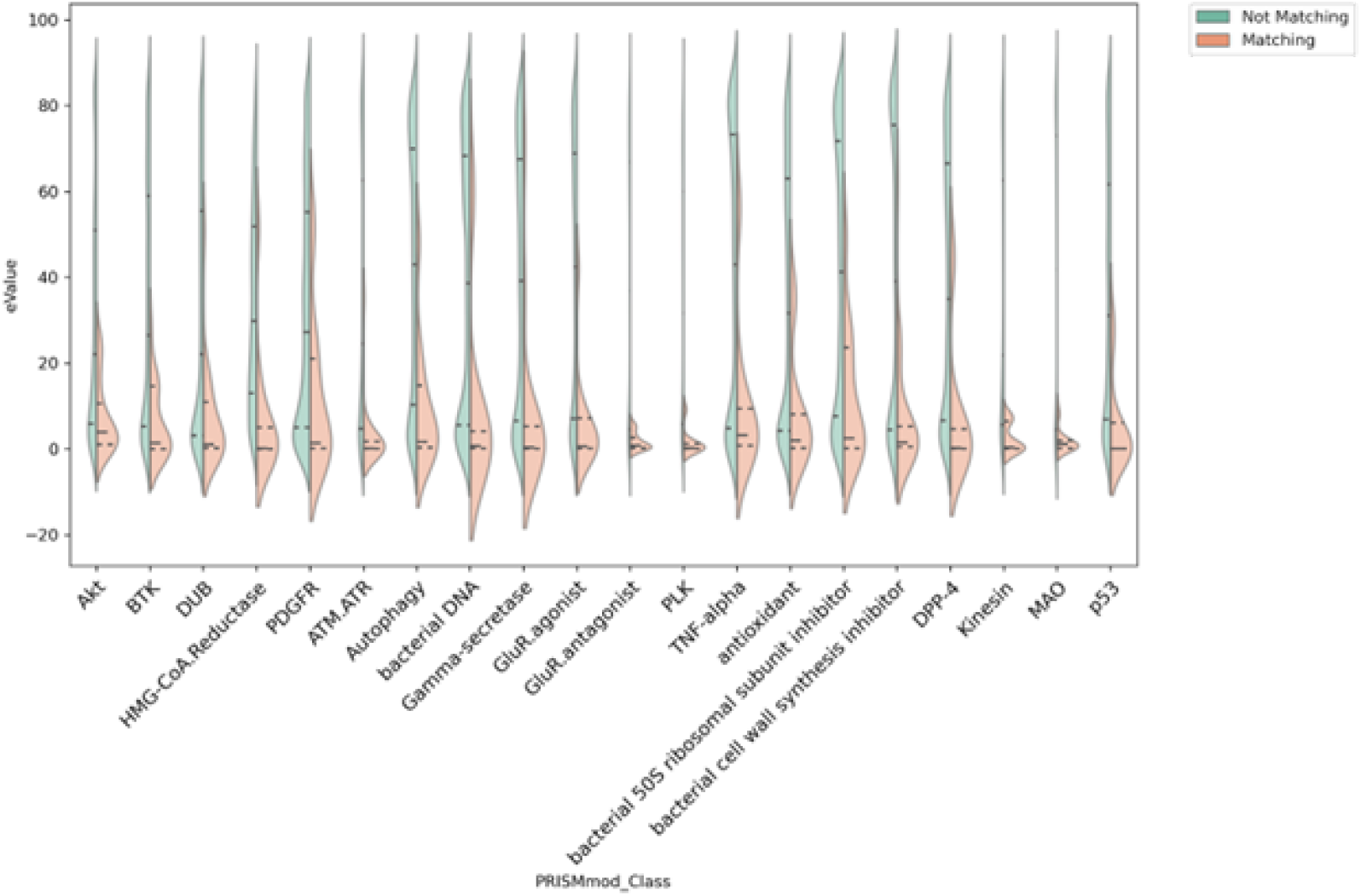
Distribution of E-values calculated for a matching or mismatched annotation.

**Fig. 7.**
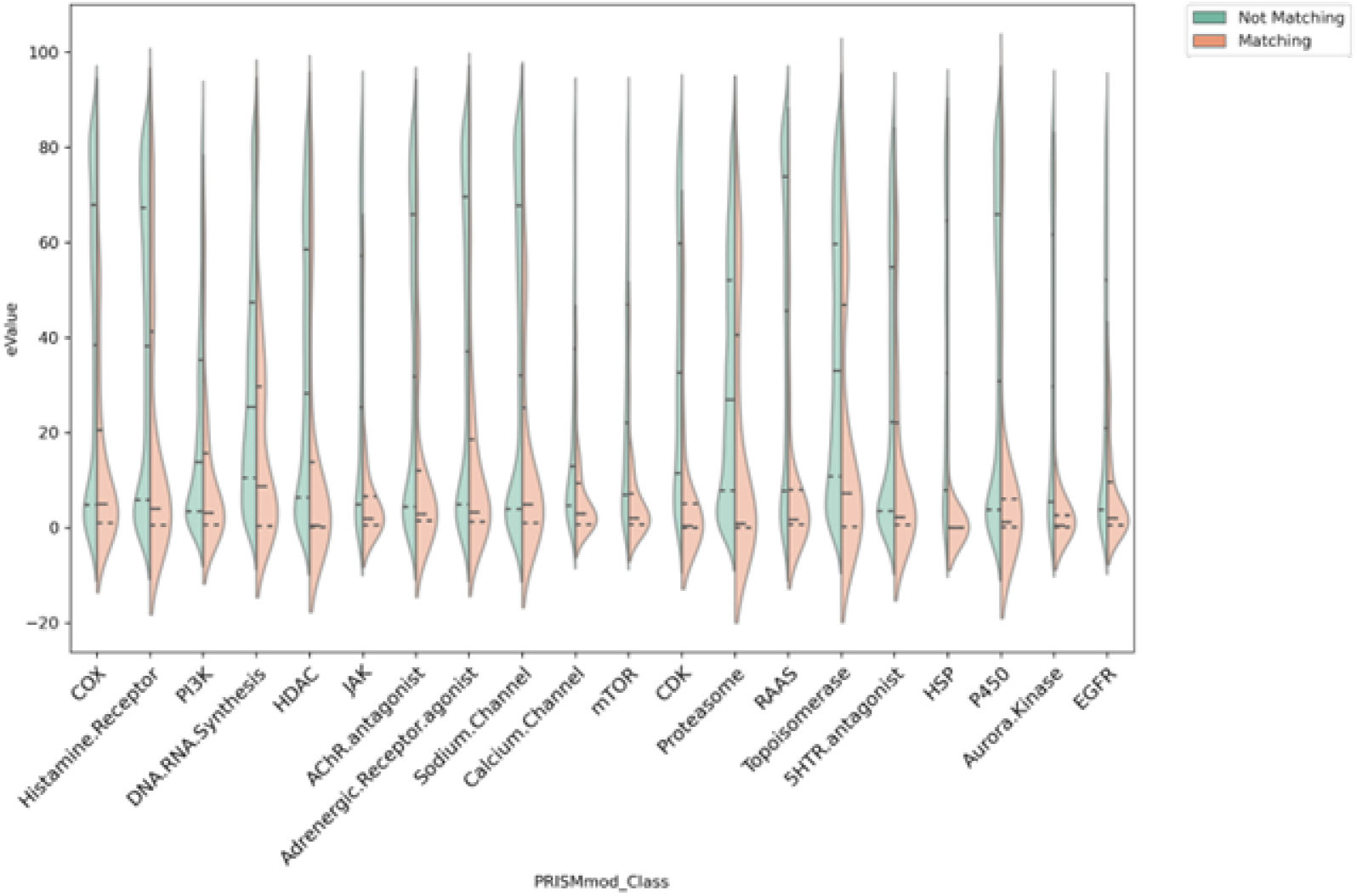
Distribution of E-values calculated for a matching or mismatched annotation.

**Fig. 8.**
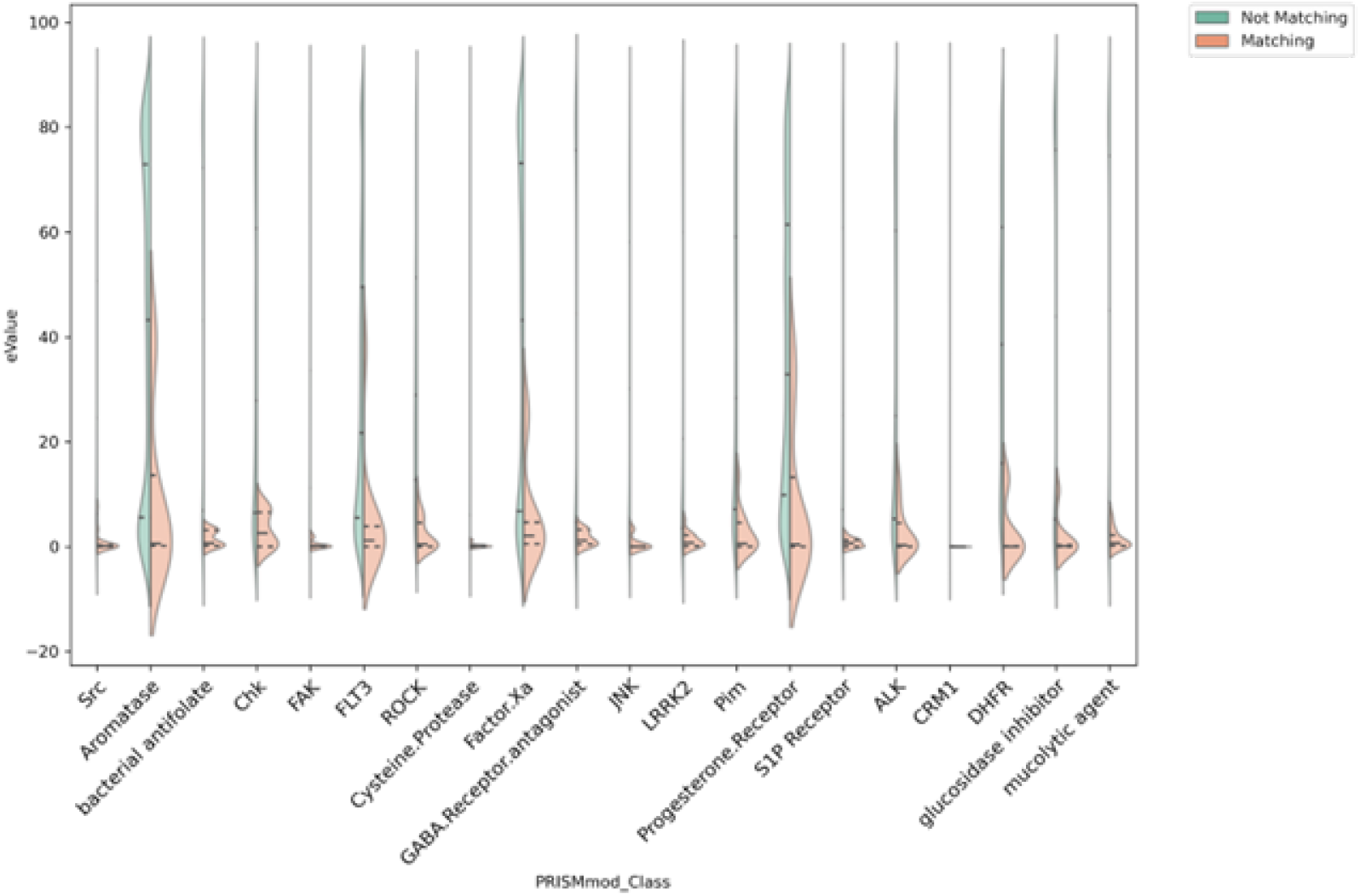
Distribution of E-values calculated for a matching or mismatched annotation.

**Fig. 9.**
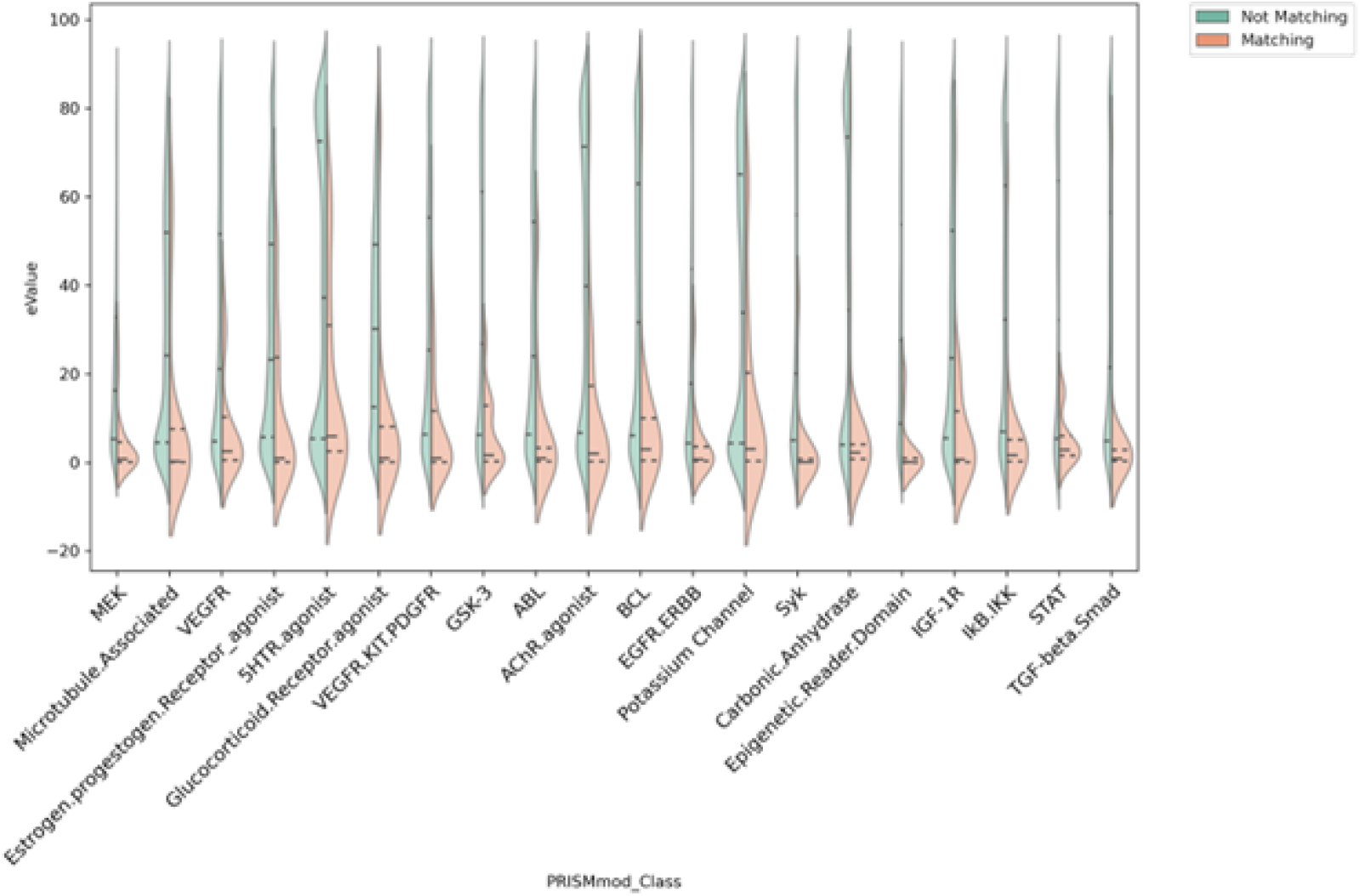
Distribution of E-values calculated for a matching or mismatched annotation.

**Fig. 10.**
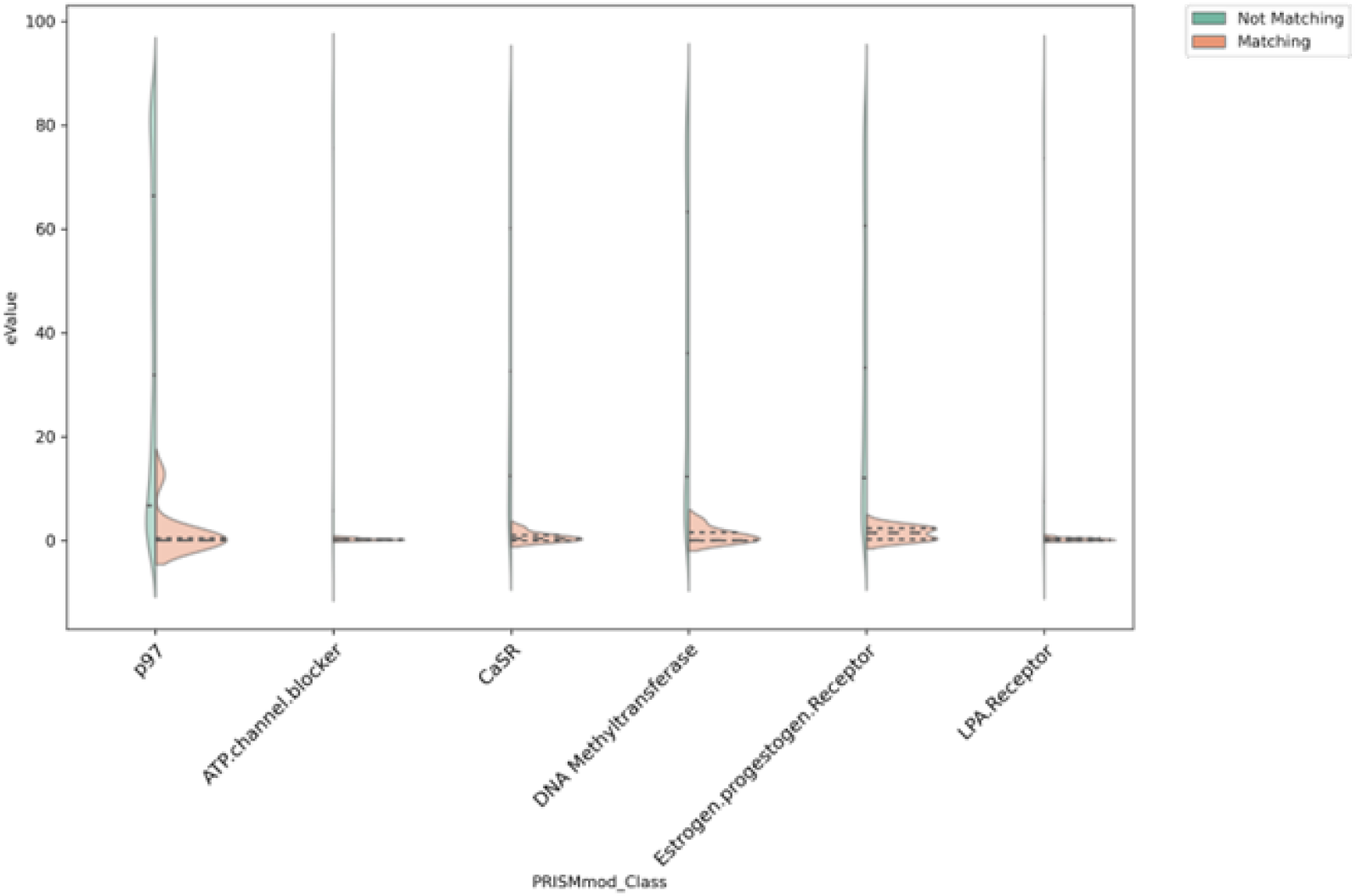
Distribution of E-values calculated for a matching or mismatched annotation.

**Fig. 11.**
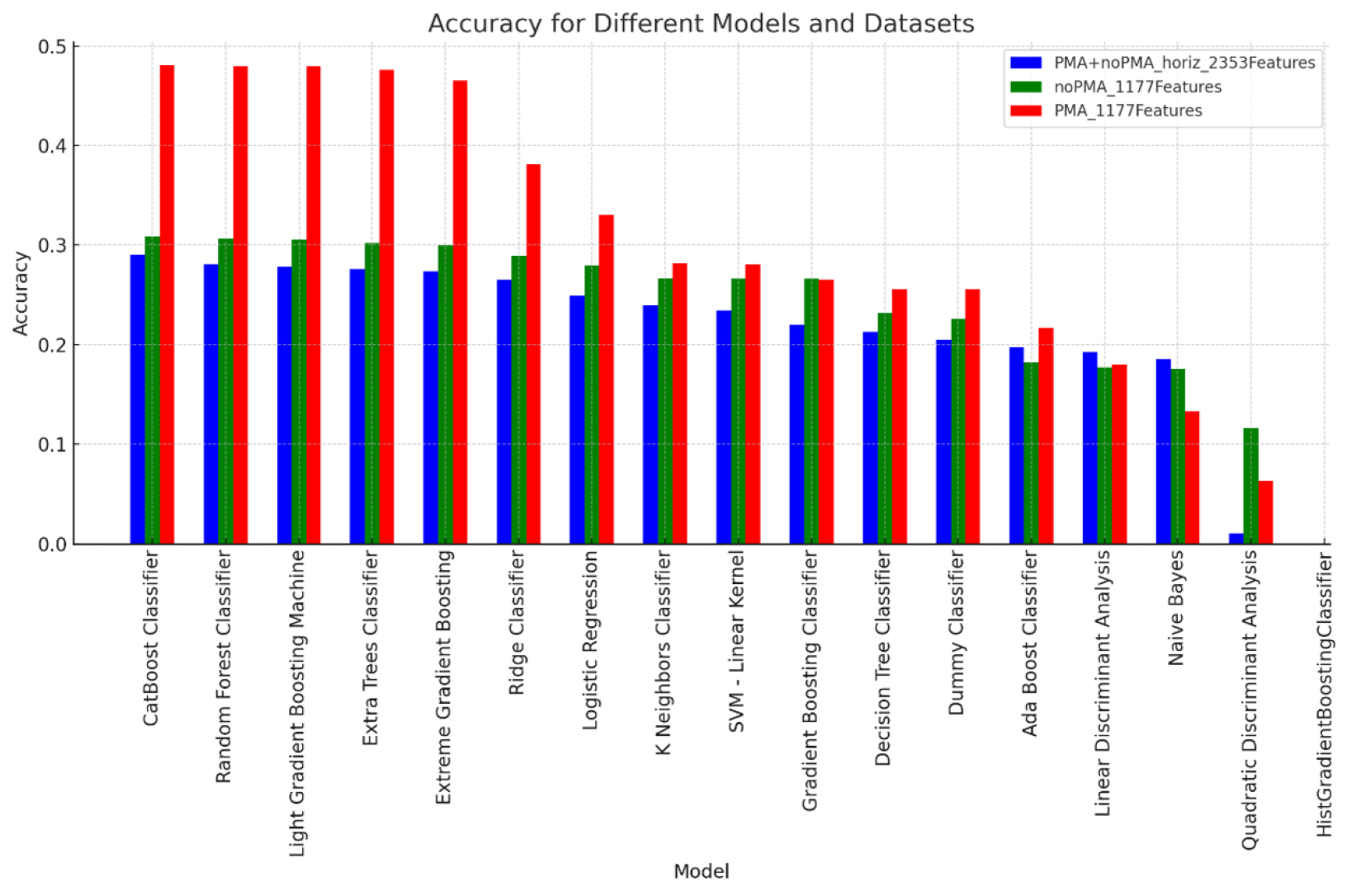
Comparison of Top Accuracy classification metric of ML models using pycaret’s default feature selection methodology (top 20% features from a fitted Light Gradient Boosting Machine). Condition datasets are treated independently and horizontally concatenated.

**Fig. 12.**
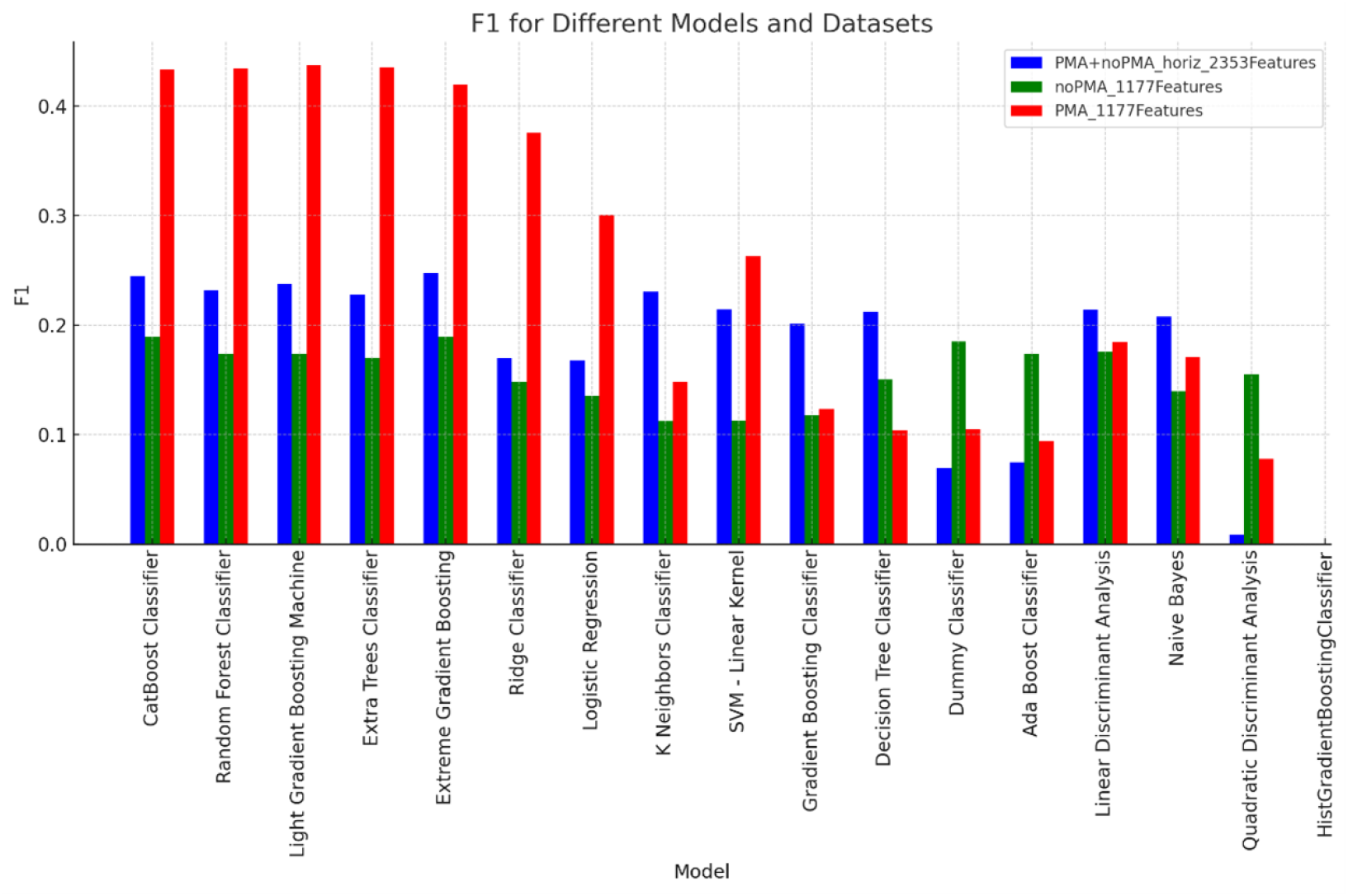
Comparison of F1 classification metric of ML models using pycaret’s default feature selection methodology (top 20% features from a fitted Light Gradient Boosting Machine). Condition datasets are treated independently and horizontally concatenated.

**Fig. 13.**
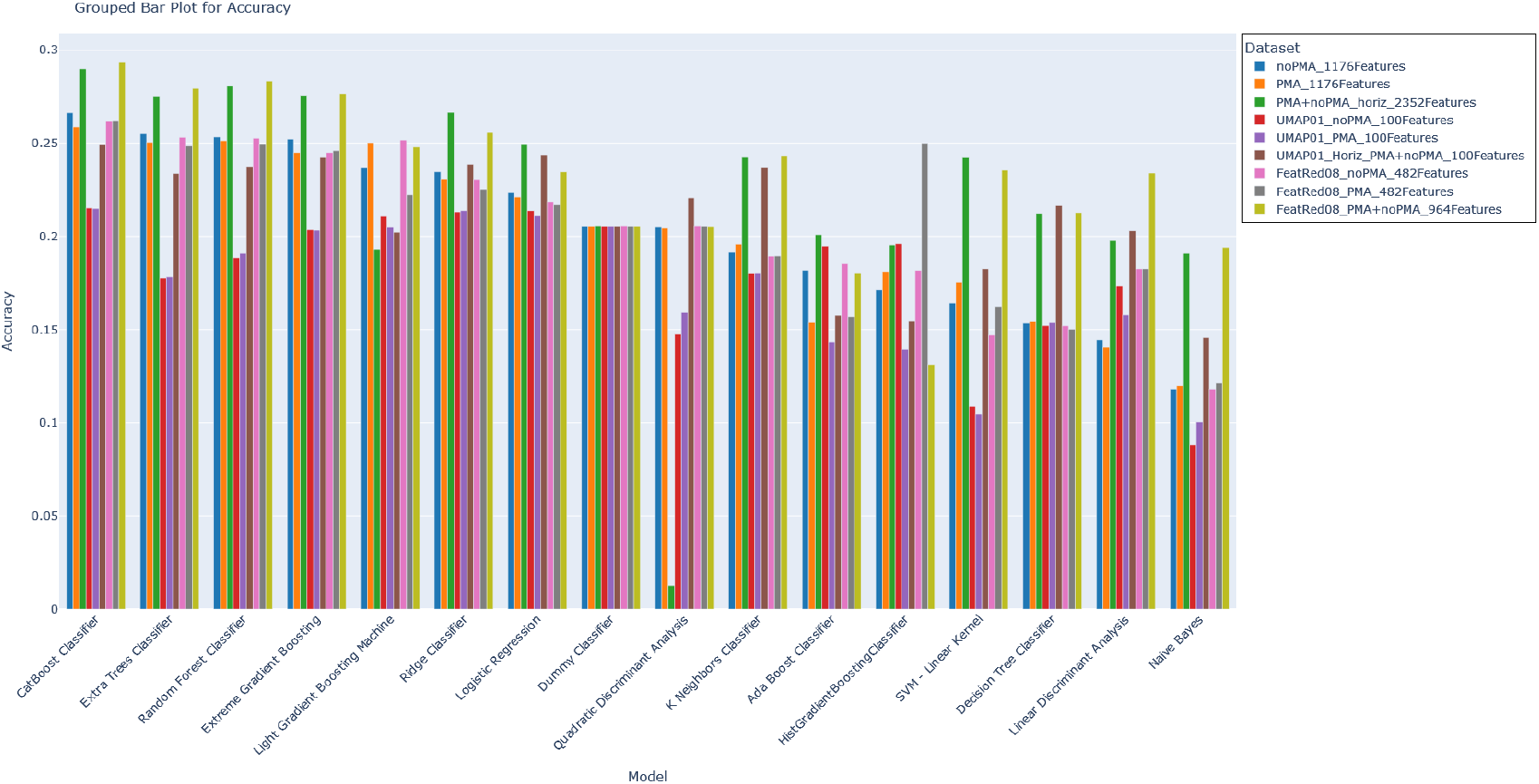
Comparison of Top 1 Accuracy classification metric of ML models using different feature reduction methodologies

**Fig. 14.**
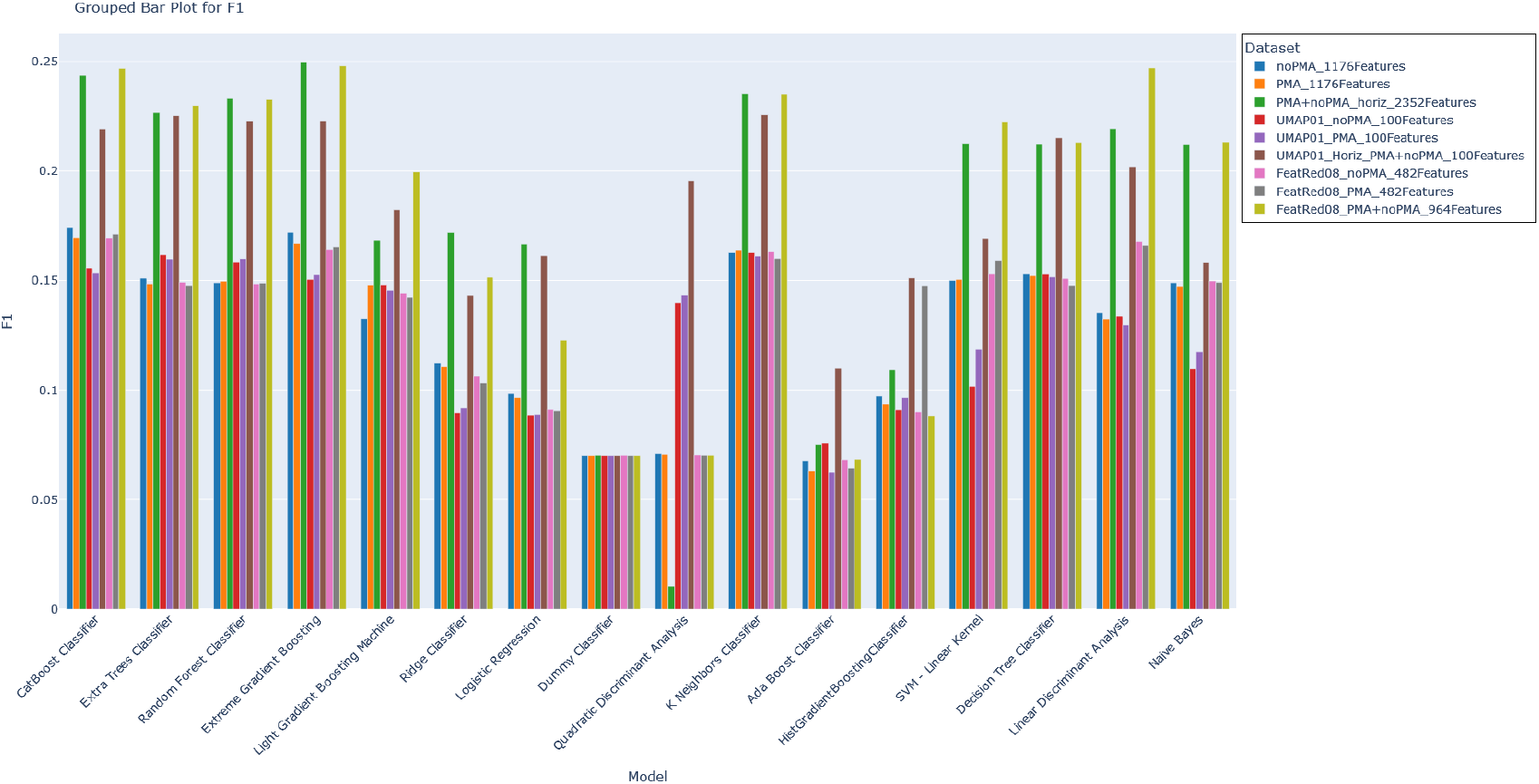
Comparison of F1 classification metric of ML models using different feature reduction methodologies

**Fig. 15.**
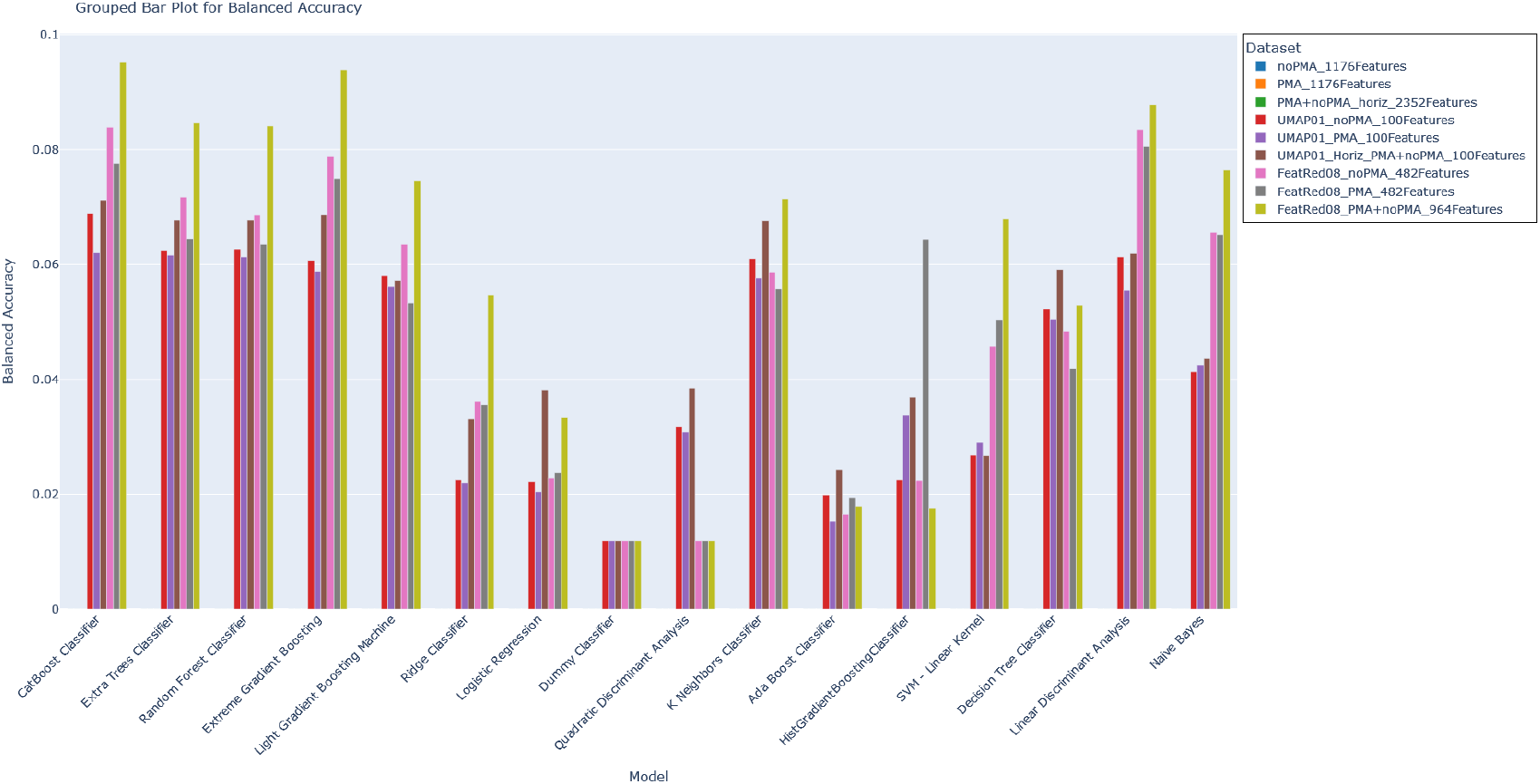
Comparison of Balanced Accuracy metric of ML models using different feature reduction methodologies

In plots 6–10, we have each class that has a significant P-Value based on the KS-test from the SelleckChem HeLa CP assay data and violin plots depicting the distribution of E-Values for the held-out testing fingerprints split based on whether the tested query was an annotated for the class or not. The 25th,50th, and 75th quartiles of the E-Values in each category are superimposed as dotted lines on the respective half-violin plots.

Breaking down the distributions by MOA class to verify that better than the median mismatch E-Value is a good way to determine a global threshold of a ‘good’ E-Value, it becomes evident that the assumption that a query fingerprint could be assigned into a single MOA annotation partition is the true culprit of the poor accuracy scores, and even ‘better than the median mismatch’ E-Value may not apply to all classes. This highlights a limitation of the procedure, such that class annotations are treated independently from each other, giving reliance on the correct annotation appearing in the ‘Best Top 5’ to increase the accuracy. This suggests that if the annotations in question were more general, such as pathway terms, this methodology might still be effective.

### 3.3 Machine Learning Classifier Validation

The A549 CP assay data on the TargetMol library was used with the pycaret autoML model testing package to determine the ‘best’ ML classifier model for this type of phenotypic screening data. Specifically, the MOA classes with at least 25 compound entries were considered, explicitly excluding the ‘Others’ class. As feature selection for the phenotypic fingerprint is very important, several iterations of reduced feature fingerprints were tested. The different feature reduction methodologies compared here are the feature reduction workflow as implemented in the High-Throughput Functional Annotation of Natural Products by Integrated Activity Profiling paperHight et al (2022) with a 0.8 threshold (FeatRed08*), UMAP embeddings learned from the full 5880 features with the two stimulation conditions concatenated horizontally (increasing feature space to 11760) or vertically (2 signatures for each compound, each coming from a stimulation treatment) (UMAP01*), and the default method by which the pycaret package performs feature selection, applying a top 20% feature cutoff with a fitted Light Gradient Boosting Machine (LightGBM) ML model. Interestingly, when each condition dataset is treated separately in model testing, the features selected with the LightGBM on the PMA condition only dataset are drastically better than the noPMA condition only dataset or the horizontally concatenated PMA+noPMA dataset 11,12. The A549 CP assay is designed to look at both stimulated and unstimulated conditions, and for future plates screened, having a common and consistent set of features to measure and report on suggests that the features selected for by pycaret should ‘know’ about the other condition as to represent features with the most dynamic and representative information given by the phenotypic fingerprint, regardless of condition assayed.

When picking an ML classifier architecture, considering the best-performing accuracy for the top result provides a good starting point for reporting the expanded classification task of the top 5 putative MOA classes. Across all the models tested, by and large the CatBoost Classifier Prokhorenkova et al (2017) performed the best across the primary metrics used for evaluation (F1 score 14, Accuracy 13, and Balanced Accuracy 15).

The CatBoost Classifier Prokhorenkova et al (2017) is a machine-learning algorithm based on gradient boosting over decision trees. It can handle numerical and categorical variables natively and robustly across various data types. CatBoost builds balanced, symmetric trees, which are generally faster to execute and can reduce overfitting. It does this using a permutation-driven approach to constructing the decision trees. Each iteration uses a different data permutation to prevent the learner from focusing too much on specific features early in the training process. Additionally, the CatBoost Classification algorithm includes an overfitting detection method. This works by stopping the training when the model starts to overfit, making the training process more efficient and potentially more effective. Designed to be efficient and scalable, CatBoost handles large datasets well, can be trained on GPU, and is faster than many other gradient-boosting libraries when working with categorical data. Despite being a complex ensemble model, CatBoost provides tools for model interpretation, such as SHAP values, which help in understanding the contribution of each feature to the model’s predictions.

Despite having a one-vs-all class-based training strategy, where the multi-class classification problem is broken down into a series of multiple binary classification problems, the ordered boosting method used in the construction of the decision trees ensures balance across all training classes such that over-fitting to specific classes does not occur early in the training process. Additionally, by keeping a probability distribution of each class for a given input, more nuanced patterns in the data are captured by considering the probabilities of all classes.

Figure 16 depicts the pipeline to obtain a trained CatBoost Classifier, and figure 17 depicts the primary metrics we are using to evaluate the ML models tested to see how the Catboost Classifier model architecture performs with an increased number of classes considered, classes with at least three representative compounds. Combining fingerprint information for both conditions, either before UMAP embedded transformation or with the feature reduction process, drastically improves the accuracy of the top 1 and F1 metrics. Unfortunately, reproducing the same results for the feature set determined by the LightGBM 20% cutoff could not be produced with the resources we applied to the problem, probably due to the large number of MOA annotations to fit in the CatBoost Classification model. It is interesting to see that while the inclusion of both PMA-stimulated and unstimulated conditions in the feature selection process reduces the classifier’s performance when trained with just the stimulated data, a classifier trained with data from both condition experiments can be more obviously improved.

**Fig. 16.**
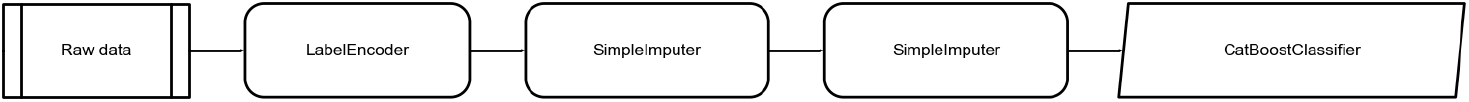
Pycaret CatBoost Model Training Pipeline

**Fig. 17.**
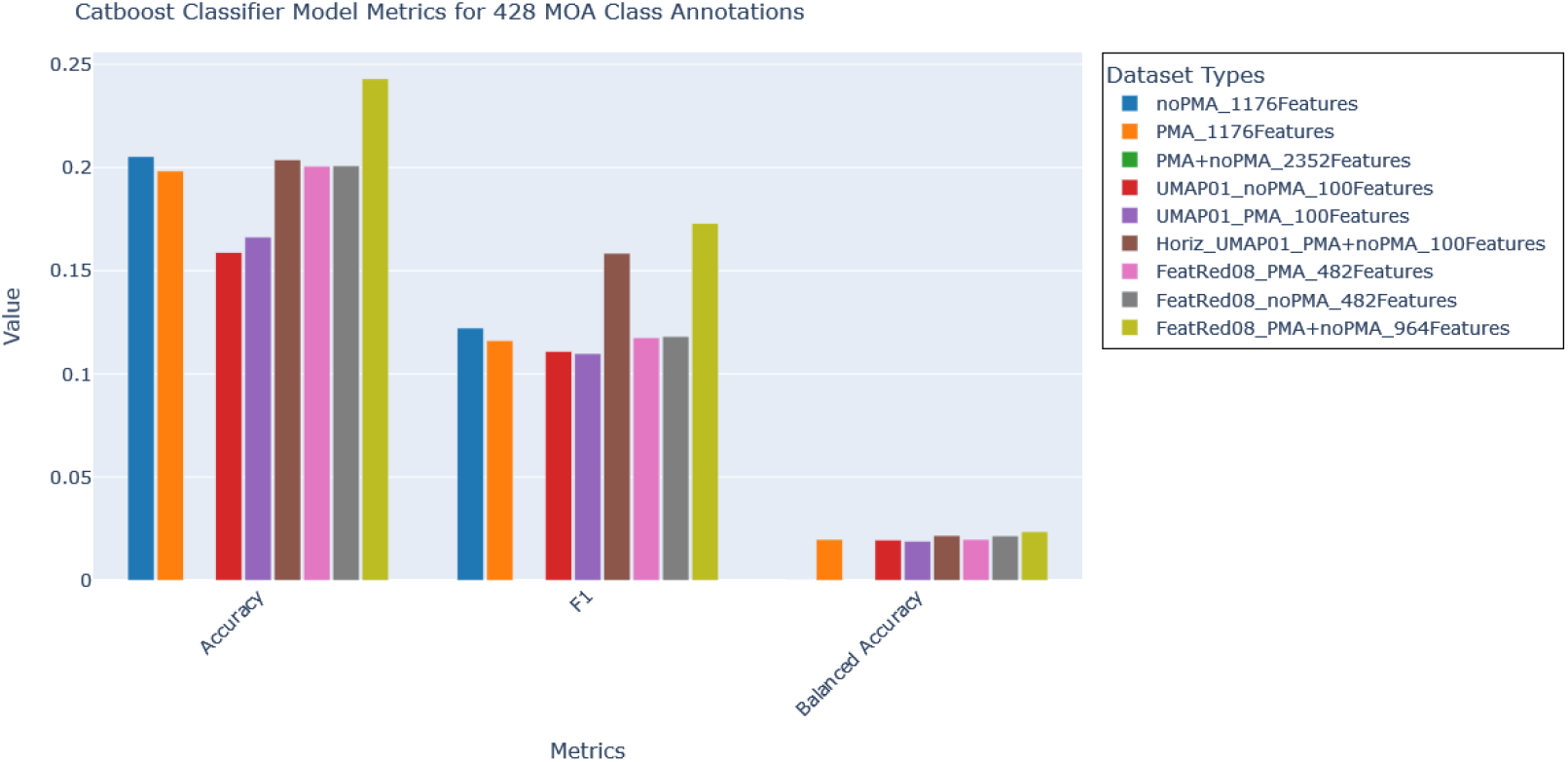
CatBoost ML Model Evaluated Metrics for training models on the data with many MOA annotations using information from the A549 CP assay data on TargetMol reference library compounds. MOA annotations considered to be trained include all classes with at least 3 compounds (regardless of experimental condition).

When we expand the classification accuracy results from the top 1 to the top 5 (figures 19, 18), we can see a significant improvement in accuracy, encompassed by the expansion of the considered MOA classes to be predicted and reported. The distributions of the prediction probabilities for each of the top 1 to the top 5 results reflect this as well (Figures 20, 21). The large jump in improved accuracy for the UMAP-trained models when expanding from matching in the top 1 to the top 2 is probably explained by the relatively low prediction probabilities for matches versus the prediction probability of the non-matches, as seen in 20.

## 4 Conclusion

MOAST addresses a fundamental challenge in natural products drug discovery: the cost-prohibitive nature of traditional screening approaches that require screening large numbers of annotated compounds alongside unknowns to determine the mechanism of action. By following the BLAST concept, MOAST allows researchers to screen just their unknown sample and compare it with an existing annotated reference set to obtain putative mechanism of action hypotheses expediently.

The tool explores two complementary approaches that address different aspects of MOA determination. The KDE integration method, inspired by BLAST, provides statistical rigor through E-value calculations and treats each MOA class independently. However, this approach implies mutual exclusivity between tested annotations, making ranking results challenging. The CatBoost classifier approach overcomes this limitation by learning about annotations in a unified framework, providing ranked predictions with prediction probabilities that enable the establishment of a global threshold for trustworthy results.

Validation results demonstrate MOAST’s effectiveness across different experimental conditions. The KDE integration method achieved approximately 22% accuracy when considering the top 5 predictions among ∼300 classes, with significant improvements when limiting to classes with sufficient members for statistical testing. The CatBoost classifier showed superior performance, with balanced accuracy reaching almost 10% for classes with at least 25 compound representatives—significantly better than the ∼3% typically reported in the literature for similar phenotypic screening approaches.

The tool’s performance across multiple feature reduction methodologies—including data-driven collinearity reduction, UMAP embeddings, and PCA—demonstrates its adaptability to different data preprocessing approaches. The PMA-stimulated condition data performed significantly better than unstimulated or concatenated datasets, suggesting that PMA stimulation effectively exposes more biological activity. The concatenated dataset with collinearity-reduced features performed comparably to LightGBM-selected features while using approximately half as many features.

MOAST’s practical utility extends beyond academic research to industrial drug discovery, where rapid MOA hypothesis generation can accelerate lead compound development and reduce costs associated with target identification. The tool’s database query model allows for rapid assessment of phenotypic fingerprint similarities without requiring extensive model training or large labeled datasets, making it accessible to researchers with limited computational resources or smaller compound libraries.

Future development of MOAST will focus on expanding the reference database with additional compound libraries and cell lines, integrating with other phenotypic screening platforms beyond cytological profiling, and developing user-friendly interfaces for broader accessibility. The modular design allows for easy adaptation to different assay types and feature sets, making it a versatile tool for the drug discovery community as phenotypic screening continues to evolve with new imaging technologies and multi-omics approaches.

In conclusion, MOAST represents a significant step forward in making MOA determination accessible to a broader range of researchers in natural products drug discovery. By combining statistical rigor with practical usability, the tool addresses key limitations of current approaches while maintaining the flexibility needed for diverse research applications. The establishment of a prediction probability threshold for trustworthy results, combined with the significant improvement in accuracy when expanding from top 1 to top 5 predictions, validates MOAST’s design philosophy of providing ranked putative annotations that acknowledge the complex, multifaceted nature of compound action.

## Supplementary information

**Fig. 18.**
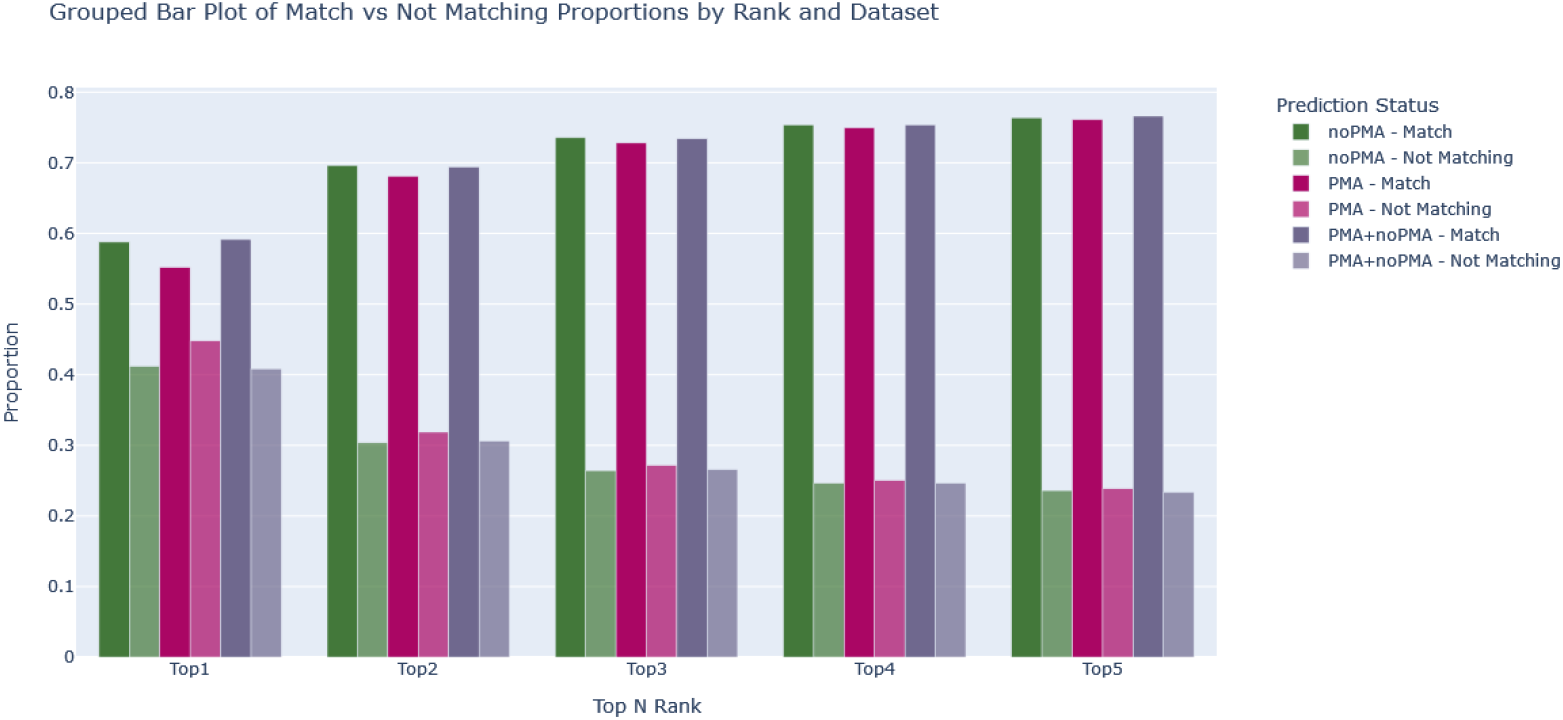
Proportions of matching or not matching predictions for CatBoost Classifier model trained on each UMAP feature embedded fingerprint dataset when considering the top 1-5 predictions

**Fig. 19.**
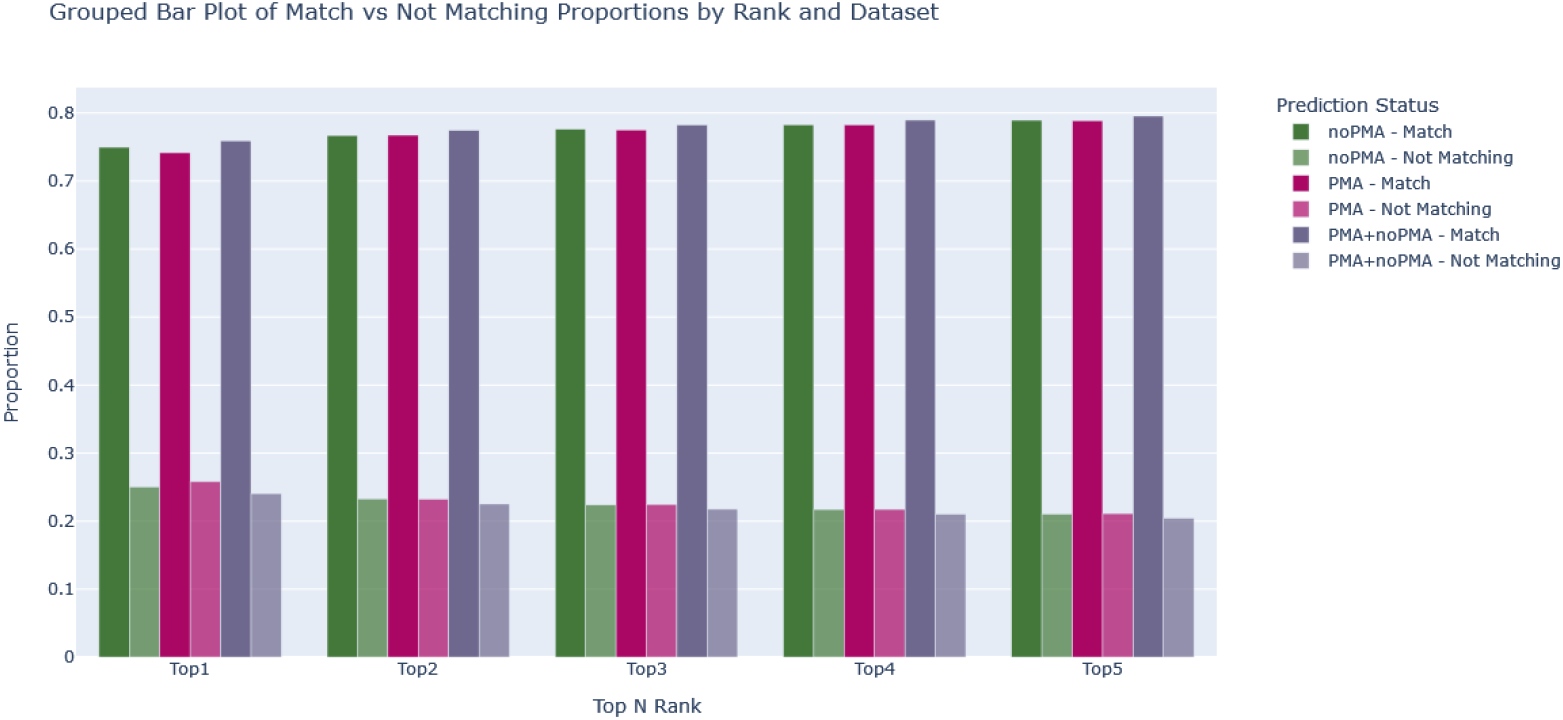
Proportions of matching or not matching predictions for CatBoost Classifier model trained on each Feature Reduction processed fingerprint dataset when considering the top 1-5 predictions

## Funding

Work funded by The National Institutes of Health (NIH) National Center for Complementary and Integrative Health (NCCIH) and Office of Dietary Supplements (ODS), NIH-U41AT008718

**Fig. 20.**
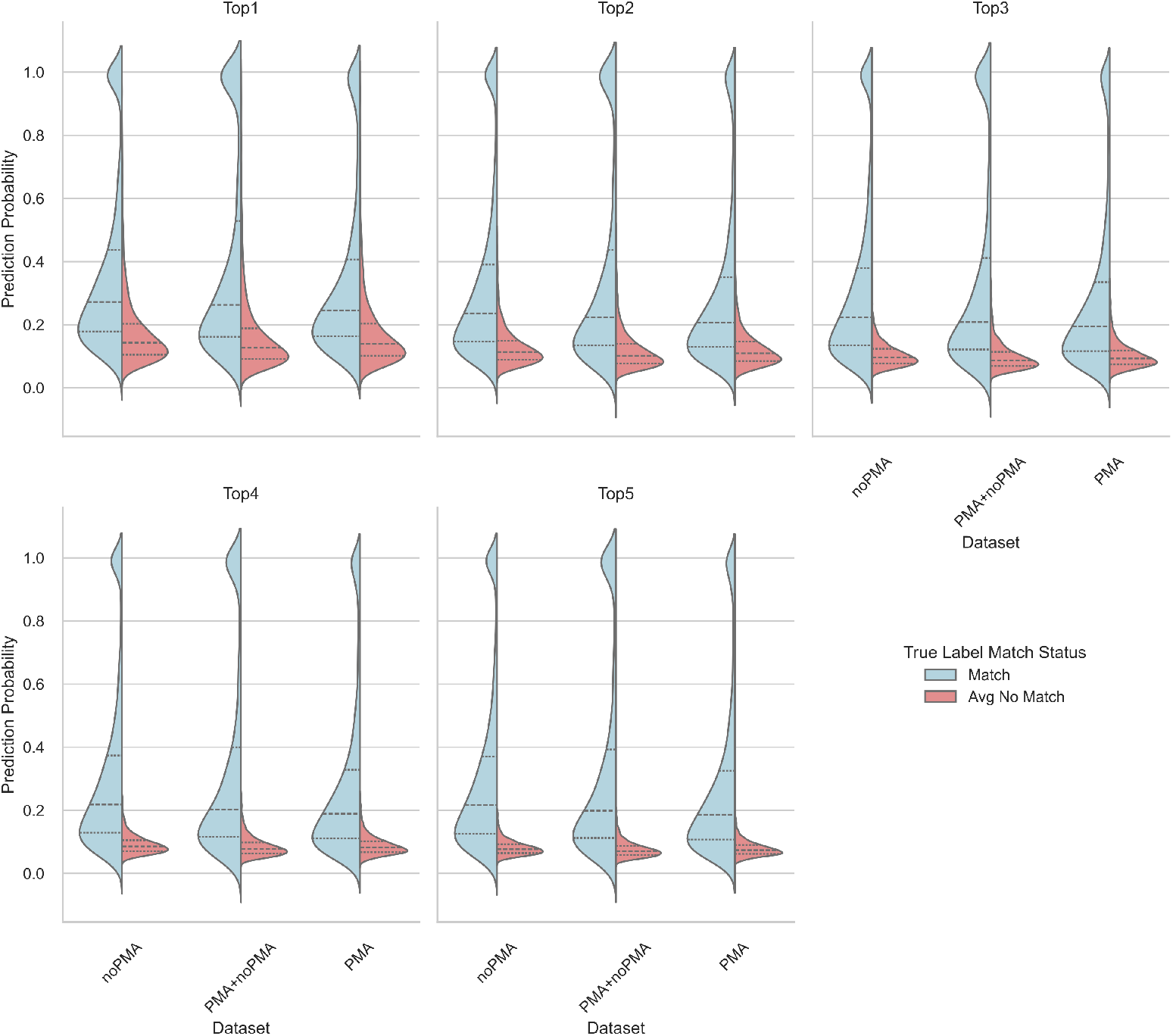
Distributions of prediction probabilities of the CatBoost Classifier models (trained on each UMAP embedded fingerprint dataset) if the annotation matches the top 1-5 rank of the predictions, as well as the average prediction probabilities if an annotation match is not found among the top 1-5 predictions.

## Data Availability

The cytological profiling data used in this study is available as associated with the High-Throughput Functional Annotation of Natural Products by Integrated Activity Profiling paper Hight et al (2022) and Cell Painting in activated cells illuminates phenotypic dark space and uncovers novel drug mechanisms of action paper Zietek et al (2025) or from the corresponding author upon reasonable request.

## Code Availability

The MOAST tool and associated code is available at https://github.com/alohith/MOAST.

**Fig. 21.**
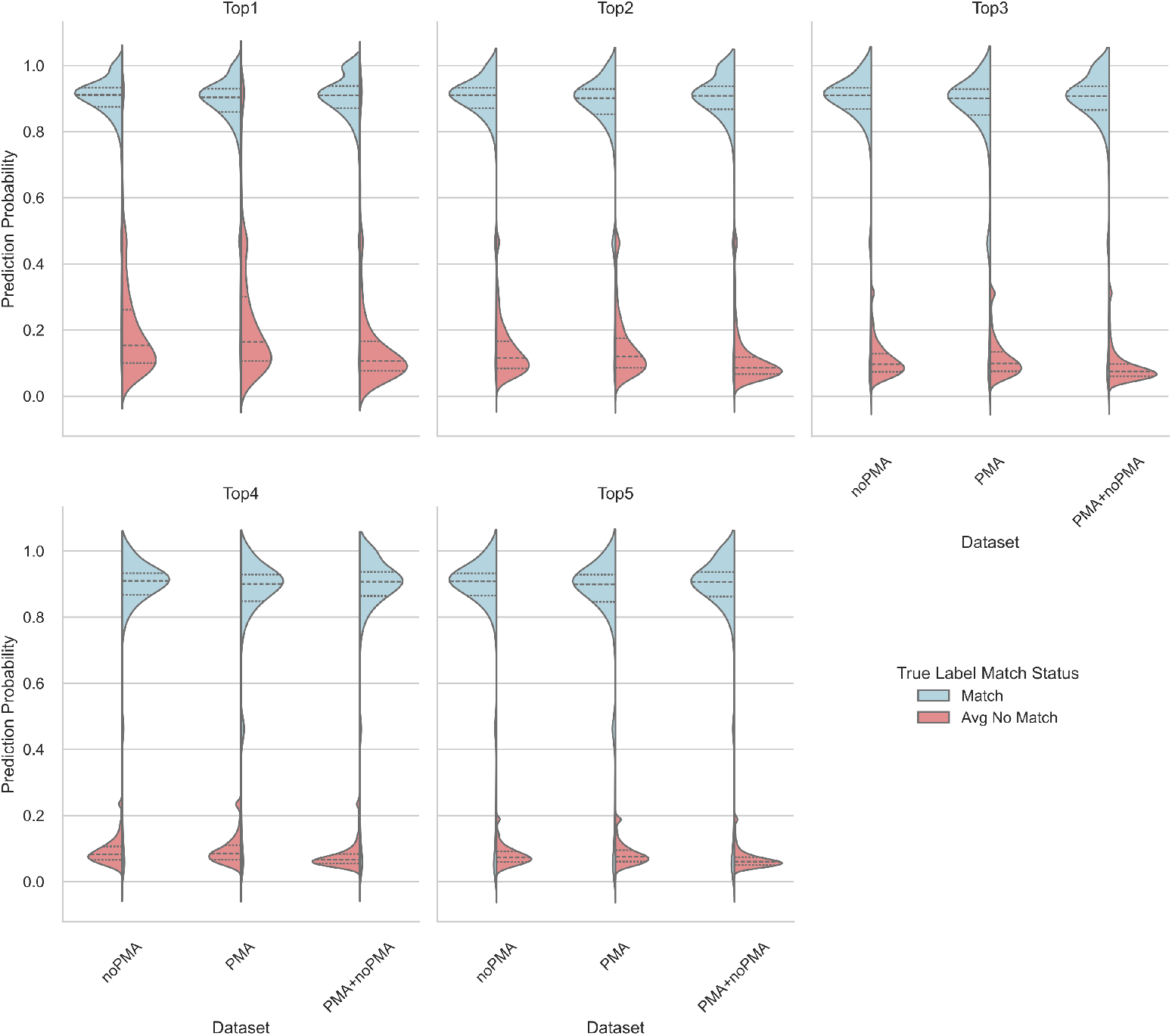
Distributions of prediction probabilities of the CatBoost Classifier models (trained on each Feature Reduction processed fingerprint dataset) if the annotation matches the top 1-5 rank of the predictions, as well as the average prediction probabilities if an annotation match is not found among the top 1-5 predictions.

**Fig. 22.**
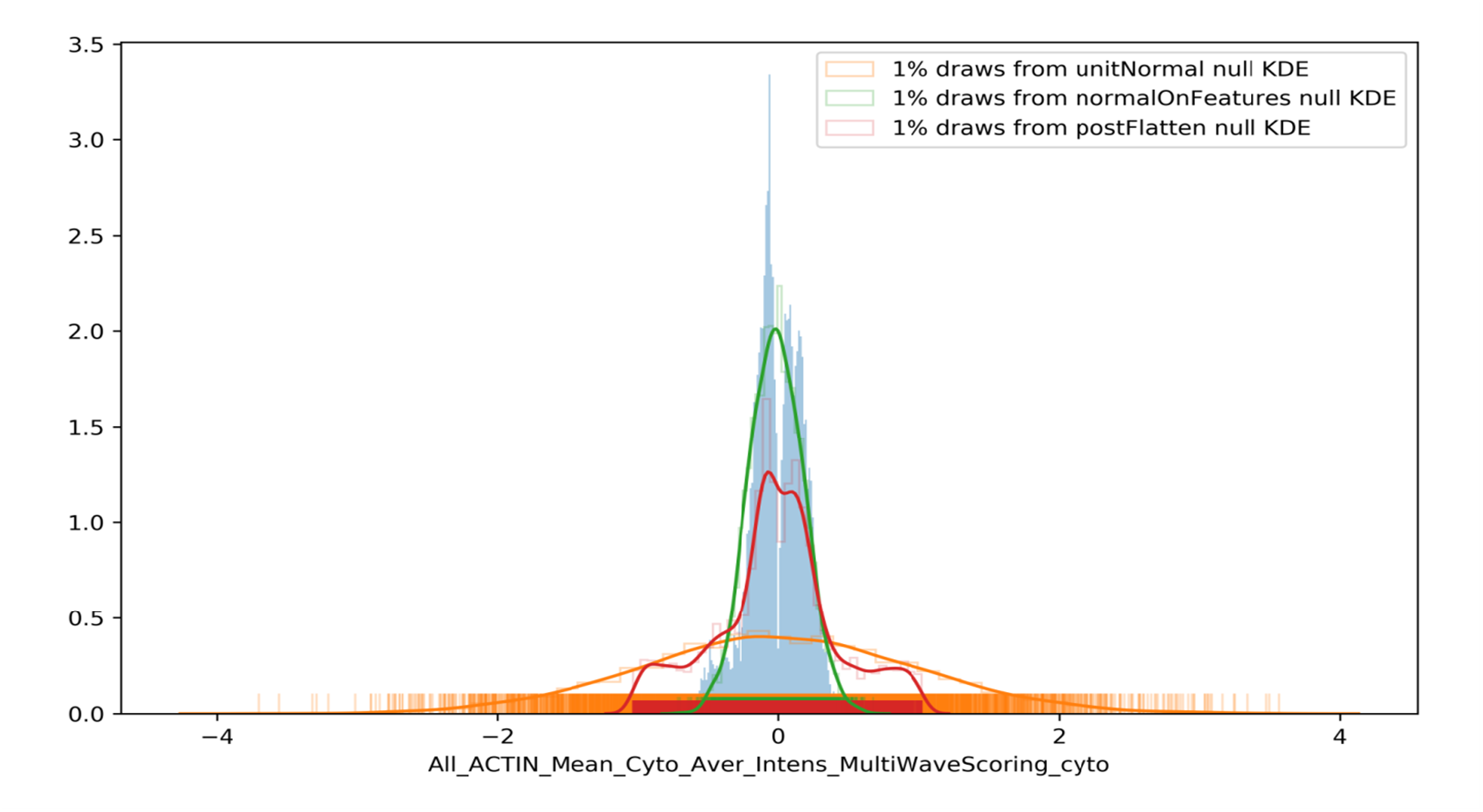
Example of Feature Null Distributions drawn on top of a histogram of the historical HistDiff scores for the Mean of all the Cytoplasmic Average Intensities in the Actin channel (‘All ACTIN Mean Cyto Aver Intens MultiWaveScoring cyto’) feature for the creation of artificial fingerprints.

## Notes

https://drive.google.com/drive/folders/1COMpRT6sG1v4k6X_JVN1tuQwmQjzpD0G?usp=sharing

### Competing Interest Statement

The authors have declared no competing interest.

